# RankCompV3: a differential expression analysis algorithm based on relative expression orderings and applications in single-cell RNA transcriptomics

**DOI:** 10.1101/2023.11.28.569110

**Authors:** Jing Yan, Qiuhong Zeng, Xianlong Wang

## Abstract

Effective identification of differentially expressed genes (DEGs) has been challenging for single-cell RNA sequencing (scRNA-seq) profiles. Many existing algorithms have high false positive rates (FPRs) and often fail to identify weak biological signals. Here, we present a novel method for identifying DEGs in scRNA-seq data called RankCompV3. It is based on the comparison of relative expression orderings (REOs) of gene pairs which are determined by comparing the expression levels of a pair of genes in a set of single-cell profiles. The numbers of genes with consistently higher or lower expression levels than the gene of interest are counted in two groups in comparison, respectively, and the result is tabulated in a 3×3 contingency table which is tested by McCullagh’s method to determine if the gene is dysregulated. In both simulated and real scRNA-seq data, RankCompV3 tightly controlled the FPR and demonstrated high accuracy, outperforming 11 other common single-cell DEG detection algorithms. Analysis with either regular single-cell or synthetic pseudo-bulk profiles produced highly concordant DEGs with ground-truth. In addition, RankCompV3 demonstrates higher sensitivity to weak biological signals than other methods. The algorithm was implemented using Julia and can be called in R. The source code is available at https://github.com/pathint/RankCompV3.jl.

## Introduction

High-throughput transcriptomic sequencing (RNA-seq) is a powerful tool for comprehensive expression profiling, which is essential for understanding biological and medical problems. A key step in RNA-seq analysis is the detection of differentially expressed genes (DEGs) [1,2]. However, the use of RNA-seq data often involves joint analysis of data across multiple platforms, which can introduce batch effects [3]. Batch effects are systematic differences between samples that are not due to biological variation. Many batch effect adjustment methods have been proposed, such as SVA [4] for microarray data and svaseq [5] for RNA-seq data.

However, normalization of different batch datasets can also distort true biological signals [6], especially when the samples are not evenly distributed between different batches. It can distort true biological signals and lead to high false positive rates (FPRs) for DEGs. Therefore, it is important to carefully consider the normalization step when analyzing RNA-seq data. The relative expression orderings (REOs) of gene pairs within a profile are often stable in samples of the same phenotype, but differ significantly in different phenotypes. This can be used to construct biomarkers and identify differentially expressed genes (DEGs). Our lab previously developed two versions of an algorithm based on REOs, RankComp [7] and RankCompV2 [8]. We successfully applied these algorithms to microarray, RNA-seq, methylation, and proteomic data [7,9-14]. RankComp can be used to identify DEGs at the population and individual levels, and it is insensitive to batch effects and normalization.

Previous versions of the algorithm used Fisher’s exact test to calculate the significance level of a 2×2 contingency table. The two rows of the table represent two groups in comparison, such as control and treatment groups, and the two columns represent the two REO outcomes: whether the expression level of the paired gene is higher or lower than that of the current gene. The test evaluates whether there is a significant correlation between the treatment conditions or phenotypes and the distribution of REO outcomes in the two groups.

RankCompV2 evaluates the stability of DEGs by including background genes updating cycles to filter out unstable DEGs. This improvement addresses the problem that upregulation or downregulation of a gene may be incorrectly indicated by its paired gene if the paired gene is dysregulated. However, both RankComp and RankCompV2 neglect the matched pair relationship of REOs in the two compared groups. As a result, REOs that are consistent in both groups are also included in the construction of contingency tables. This negligence may lead to non-differential genes being identified as DEGs, resulting in high FPRs.

McNemar’s test is a statistical test that takes into account the matched experiment design in the test of 2×2 contingency tables. McNemar-Bowker’s test extends this to the general situation with more than 2 categories. However, both of these tests do not take into account the ordered relationship of the three possible outcomes of REOs.

Another limitation of the previous two versions of RankComp is that the contingency tables do not consider the contribution of gene pairs with approximately equal expression levels (non-stable REOs based on the binomial distribution) in one group but stable REOs in the other group.

In RankCompV3, we count the frequency of all possible 9 REO outcomes in a matched pairs design. McCullagh’s test [15], which is designed for the matched designs with ordered categories, is applied to test the 3×3 contingency tables.

We implemented RankCompV3 and tested it on single-cell RNA-seq (scRNA-seq) data. Many methods have been developed for differential expression analysis of scRNA-seq data. Some were specifically developed for scRNA-seq, such as MAST [16], DEsingle [17], Wilcoxon signed-rank test [18], Monocle2 [19], SigEMD [20], and scDD [21]. Others were originally developed for microarray and bulk RNA-seq data, such as limma [22,23], edgeR [24], and DESeq2 [25], but have been found to perform well in scRNA-seq data using the pseudo-bulk analysis scheme [26]. However, the consistency of these algorithms is not high, and many algorithms have problems with insufficient sensitivity and high FPRs. Additionally, it has been found that algorithms specifically developed for scRNA-seq data do not necessarily perform better than those designed for bulk profiles [27-29].

We evaluated the performance of RankCompV3 on scRNA-seq datasets by comparing it with several common DEG identification algorithms using both simulated multimodal single-cell datasets and real datasets. Our analysis results showed that RankCompV3 performed well on these datasets and was sensitive to weak biological signals. Additionally, we obtained concordant DEGs between the analysis comparing the single-cell profiles directly and the pseudo-bulk analysis, in which single-cell profiles were aggregated randomly into pseudo-bulk profiles [26].

## Methods and Materials

### The RankCompV3 algorithm

The RankCompV3 method for identifying differentially expressed genes (DEGs) in either bulk or scRNA-seq data consists of the following steps:

1. **Identification of significantly stable REOs**: For each gene pair (*a*, *b*), the observed REO outcome is either *a* > *b* or *a* < *b* in a single-cell profile or a bulk profile, depending on which gene has the higher expression level. If the two genes have the equal expression levels within the measurement uncertainty, the REO is randomly assigned to either *a* > *b* or *a* < *b* with equal probability. The binomial distribution model is then used to test if the observed REOs are stable across all the profiles in each group, respectively. The null hypothesis is that the two genes have the same expression levels. This means that the probability of observing each REO outcome (*a* > *b* or *a* < *b*) is equal, *p*_0_ = 0.5. The *P* value is the probability of observing the major REO outcome in *m* or more profiles out of a total of *n* profiles by chance (where ‘major’ means *m* > *n*/2) in one group, 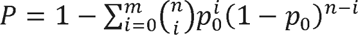. If the *P* value is less than the preset threshold, *e.g.*, *α* = 0.01, the major REO outcome is considered to be significantly stable, *a* > *b* or *a* < *b*. Otherwise, it is considered that the REO is not stable, denoted as *a* ∼ *b*.
2. **Construction of contingency tables of stable REOs**: For each gene, the contingency tables of REO counts are constructed. The contingency tables summarize the comparison results of the gene with reference genes in two groups, *e.g.*, control and treatment groups (Table 1). The diagonal elements of the contingency tables are the numbers of REOs which are consistent in the two groups, while the off-diagonal elements are the numbers of inconsistent REOs in the two groups. Lower triangular elements support that *a* is down-regulated in the treatment group compared with the control group, while the upper triangular elements support that *a* is up-regulated.
3. **Significance test of contingency tables**: McCullagh’s method is applied to test the significance of the contingency tables. The method applies a logistic model for matched comparisons with ordered categorical data. The null hypothesis *H* _0_ is that the distribution of the REO outcomes has no association with the grouping and the contingency table should be symmetric. The Benjamini-Hochberg (BH) method is used to adjust *P* values for multiple comparisons. If the adjusted *P* value < 0.05, the null hypothesis *H* _0_ is rejected, and the gene is considered as a candidate DEG.
4. **Iteration:** Initially, each gene is compared with a list of housekeeping genes to construct the contingency tables. If such gene list is not provided, all the genes are used as the reference genes. After the first cycle, all non-differentially expressed genes from the previous cycle are used as the reference list to construct new contingency tables. The cycle ends when the list of candidate DEGs does not change any more.

**Table 1.**
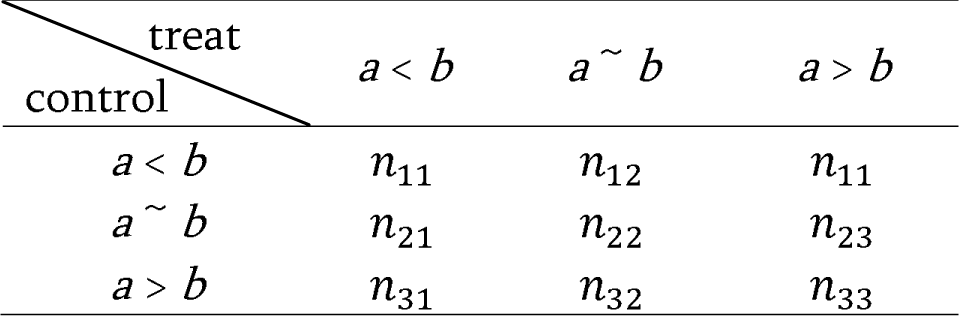
3 x 3 contingency table of REO counts for one gene (*a*). The table shows the number of REOs for gene *a* that form either a significantly stable REO or a nonstable REO with a reference gene *b* in the two groups, *e.g.*, control and treatment. The row and column headers indicate the different types of REOs. Each cell sums the number of REOs forming 9 possible paired outcomes in the two groups.

The overall workflow of the RankCompV3 method is shown in Figure 1.

**Figure 1.**
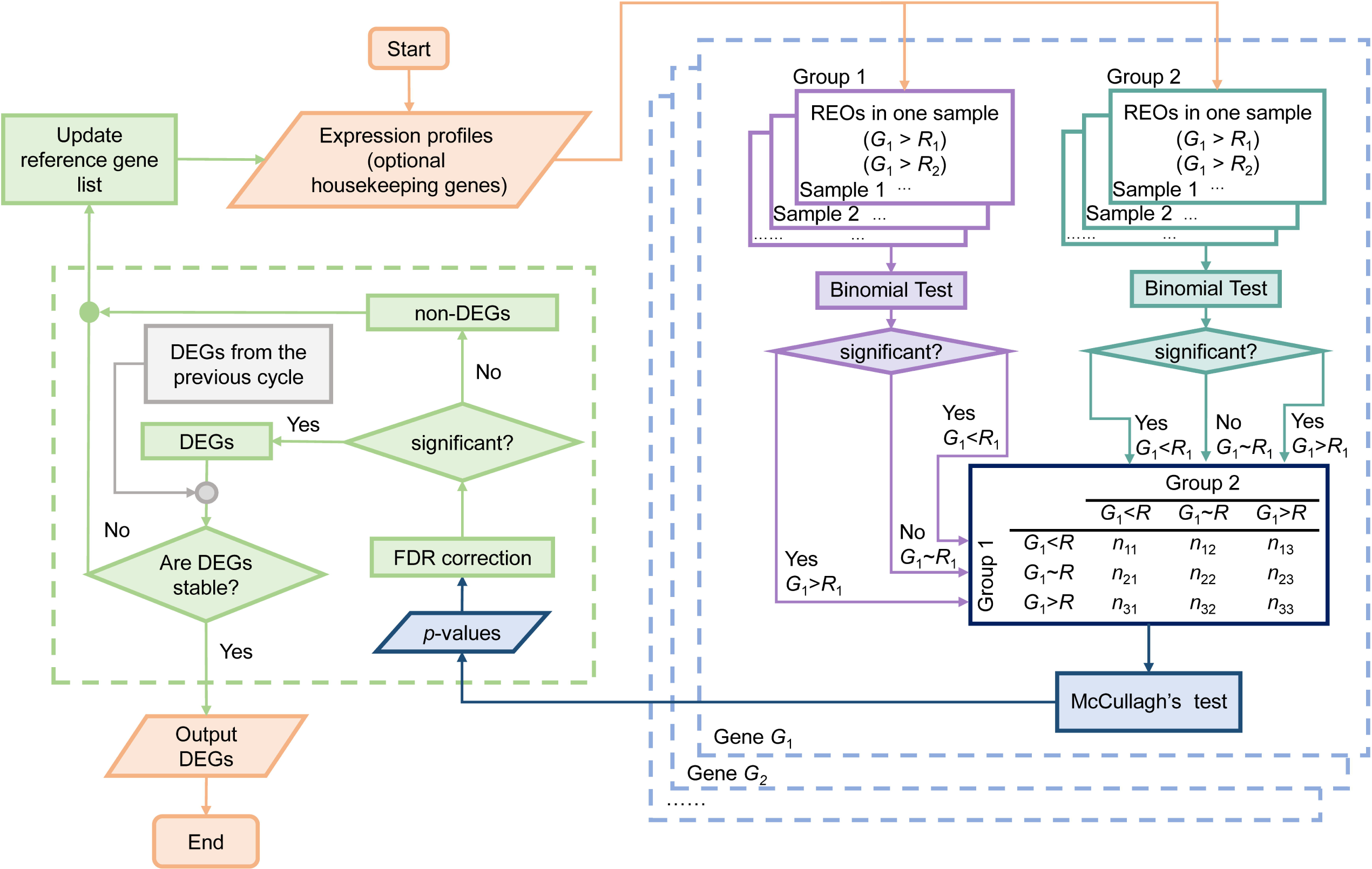
Flowchart of the RankCompV3 method. The RankCompV3 method is a three-step process for identifying differentially expressed genes (DEGs) in RNA transcriptomics. The first step is to calculate the relative expression orderings (REOs) of gene pairs in the two groups. The second step is to test the contingency tables of REOs using McCullagh’s method. The third step is to filter the DEGs using a significance threshold.

### Other DEG identification methods

We compared the performance of RankCompV3 with the performance of 11 other methods, including 7 independent methods, limma, DESeq2, edgeR, DEsingle, SigEMD, Monocle2 and scDD, and 4 methods implemented in the Seurat package, Wilcoxon rank sum test (“wilcox”), likelihood-ratio test (“bimod”), logistic regression (“LR”) and MAST. The methods were compared using a simulated scRNA-seq dataset and real scRNA-seq datasets.

### Performance evaluation

The performance of RankCompV3 was evaluated using the following metrics: true positive rate (TPR), true negative rate (TNR), false positive rate (FPR), false negative rate (FNR), precision, accuracy, AUC (area under the receiver operating characteristic curve) [41] and AUCC (area under the concordance curve) [26]. ROC curve and AUC calculation were performed with the R package of pROC.

AUCC is a more stringent metric than AUC that is used to evaluate the concordance between single-cell or pseudo-bulk DEGs and bulk ground truth.

### Benchmark dataset

We used the ground-truth scRNA-seq dataset collected in Squair et al.’s study [26] to compare the performance of our method and to the benchmark results in that study. The dataset was compiled from four studies [26,30-33] in which matched bulk and scRNA-seq were carried out on the same population of purified cells under the same conditions, as detailed in Supplementary Table 1. The bulk results were considered as the ground-truth.

### Simulation dataset

The simulation dataset was generated using the scDD package (“simulateSet” function). The synthetic dataset scDatEx provided by the author was taken as the starting point. The package models single-cell expression profiles of cells under two conditions using Bayesian mixture models to accommodate the main characteristics of scRNA-seq data, including heterogeneity, multimodality, and sparsity (a large number of 0 counts).

The profiles were generated for 75 cells and 20,000 genes under each condition, including 2,000 DEGs and 18,000 non-DEGs. The 2,000 DEGs were equally distributed among four modes: DE (differential expression of unimodal genes): The expression levels of these genes are different between the two conditions, and the genes are unimodal (i.e., they belong to a single cluster); DP (differential proportion for multimodal genes): The expression levels of these genes are different between the two conditions, but the genes are multimodal (i.e., they belong to multiple clusters); DM (differential modality genes): The expression levels of these genes are different between the two conditions, and the genes change modality (i.e., they move from one cluster to another); DB (both differential modality and different component genes): The expression levels of these genes are different between the two conditions, and the genes change modality and also change the cluster they belong to. Among the 18,000 non-DEGs, one half are EE (equivalent expression for unimodal) genes and the other half are EP (equivalent proportion for multimodal) genes.

All simulation parameters were set to default values, and the resulting data were rounded to the nearest integers. The specific types and distributions of the simulated data are shown in Figure S1.

### Real datasets

We used public datasets downloaded from the Gene Expression Comprehensive Database (GEO, http://www.ncbi.nlm.nih.gov/geo/). The datasets included bulk RNA sequencing (RNA-seq) dataset (GSE82158) and single-cell RNA sequencing (scRNA-seq) datasets (GSE54695, GSE29087 and GSE59114) measured on different platforms.

For data preprocessing, we removed the genes that were not expressed in most of the cells and the cells expressing very few genes.

#### Negative test dataset

We used 80-cell profiles from the GSE54695 dataset provided by Grün D et al. [34] as a negative test dataset to evaluate FPR. The profiles were obtained by lysing frozen mouse embryonic stem cells (mESCs) under the same culture conditions and measured with the CEL-seq technique. According to Wang et al.’s study [35], the top 7277 genes were retained for analysis with the highest number of nonzero expression in all cells. We randomly divided 80 profiles into two subsets with 40 in each. Since all the profiles were generated under the same condition, there should be no DEGs between the two subsets. The randomized experiments were repeated 10 times and the average FPR was calculated [36].

#### Positive test dataset

The positive test dataset (GSE29087) of scRNA-seq were provided by Islam et al. [37], consisting of 22936 genes from 48 mouse ES cells and 44 mouse embryonic fibroblasts. We used the top 1000 DEGs as the gold standard DEGs between the two cell types, which were verified by the real-time quantitative reverse transcription PCR (RT-qPCR) experiments [27,38].

#### Influence of sample size

The scRNA-seq dataset (GSE59114) provided by Kowalczyk et al. [39] were used to evaluate the influence of sample size (number of profiles in a group) to the performance. The dataset consisted of the profiles of 89 long-term hematopoietic stem cells (LTHSCs) from young mice (2-3 months) and 135 LTHSCs from old mice (20 months). We randomly sampled the profiles of two conditions into subsets with sizes of 10, 30, 50, and 70, respectively. The DEGs were identified between the subsets with the same sample size. DEGs identified by the same algorithm in the entire dataset were used as the gold standard. The random sampling experiments were repeated 10 times for each sample size and the average performance indices were calculated as the final result.

#### Weak-signal test dataset

To test the performance of our algorithm on weak-signal detection, we used the GSE82158 dataset provided by Misharin AV et al. [40]. We compared the profiles of 4 monocyte-derived alveolar macrophages (Mo-AMs) and 4 tissue-resident macrophages (TR-AMs) from mice treated with bleomycin for 10 months. Pathway enrichment analysis was performed on the DEGs to detect biological signals.

### Pseudo-bulk method

In addition to comparing the input single-cell profiles directly, the pseudo-bulk method is also available in the implementation of RankCompV3. The single-cell profiles are randomly partitioned into a number of subsets of equal sample size, respectively, in each group. The profiles in each subset are aggregated into a single pseudo-bulk profile. Differential expression analysis is then conducted with these synthetic profiles from two groups.

For the ground-truth datasets from Squair et al.’s study [26], the pseudo-bulk profiles were generated in the same manner with the original study, where the cells were aggregated in each replicate.

### Pathway enrichment analysis

Pathway enrichment analysis was performed against the Kyoto Encyclopedia of Genes and Genomes (KEGG) database [42] with the hypergeometric distribution model [43] through the clusterProfiler package (“enrichKEGG” function). The *P*-values were corrected by the BH method for multiple tests.

## Results

### Performance on the null dataset

We tested the FPR using the null dataset of profiles of 80 cells lysed from frozen mouse embryonic stem cells under the same culture conditions, provided by Grün D et al. [34]. The dataset was randomly divided into two subsets, each with 40 profiles. In 10 randomized trials with a false discovery rate (FDR) < 0.05 as the threshold, no or very few DEGs were detected by RankCompV3, Bimod, LR, MAST, DEsingle, Wilcoxon, edgeR, limma, and scDD algorithms. On average, 43.9, 7.5, 0, 0, 4.7, 0, 0, 0, and 77.5 DEGs were detected, respectively. The Monocle2 and SigEMD algorithms detected 132.6 and 1291 DEGs, which showed relatively high FPRs. These results show that the RankCompV3 method performs well in controlling the FPR in the negative dataset.

### Performance comparison with Squair et al’s benchmark test

Squair et al. performed a benchmark test on various DEG algorithms. It included the algorithms specially designed for single-cell mode and the algorithms designed for bulk profiles. The results from bulk RNA-seq profiles of the same samples were used as the ground truth to test the performance of the algorithms in single-cell and pseudo-bulk modes. Eleven ground-truth datasets were used to evaluate the performance of RankCompV3 using the same benchmark protocol as in ref. [26]. The results are listed in Table 2. In most datasets, the concordance between the pseudo-bulk and the ground-truth is very high. The median AUCC of RankCompV3 in the pseudo-bulk mode is 0.45, which is higher than that of the best method in ref. [26] (edgeR-LRT, median AUCC = 0.38) (Figure 2). More importantly, the concordance is also high for RankCompV3 applied to single-cell profiles directly (median AUCC = 0.39), which is better than the best method for single-cell in the ref.[26] (*t*-test and Wilcoxon rank-sum test, the median AUCC is 0.24) (Figure 2).

**Figure 2.**
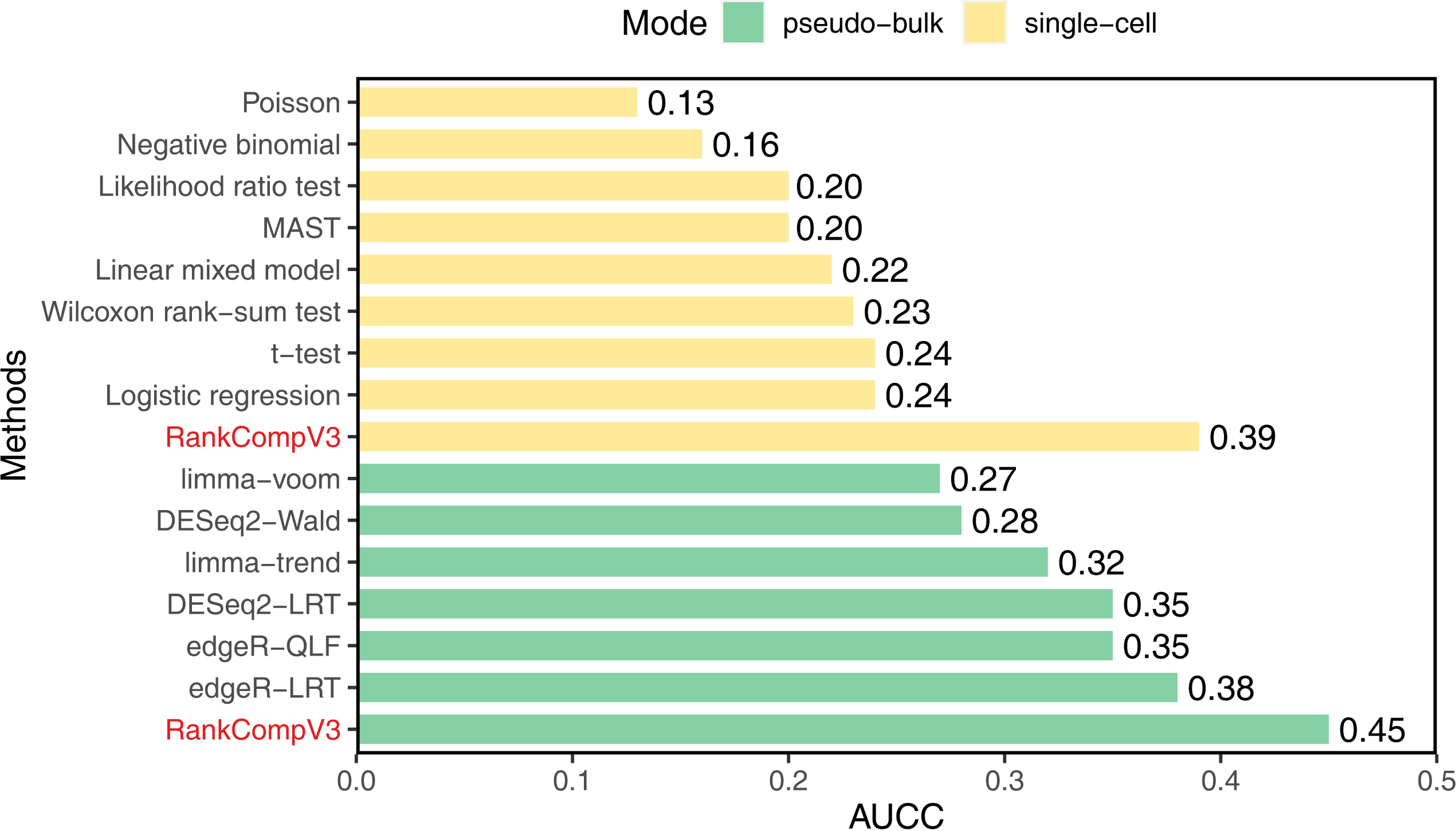
Median AUCC for RankCompV3 and 14 other DEGs recognition algorithms for pseudo-bulk analysis scheme in the eighteen ground-truth datasets from Squair et al.

**Table 2.** Detection performance of RankCompV3 and 11 other algorithms in the simulated scRNA-seq dataset (FDR < 0.05).

| Dataset | Ground-truth<sup>a</sup> |  | edgeR-LRT<sup>b</sup> |  |
| --- | --- | --- | --- | --- |
|  | Pseudo-bulk | Single-cell | Pseudo-bulk | Single-cell |
| Angelidis2019_alvmac | 0.039 | 0.028 | 0.053 | 0.049 |
| Angelidis2019_pneumo | 0.216 | 0.195 | 0.226 | 0.176 |
| CanoGamez2020_memory-iTreg | 0.382 | 0.306 | 0.186 | 0.177 |
| Hagai2018_mouse-lps | 0.742 | 0.502 | 0.457 | 0.504 |
| Hagai2018_mouse-pic | 0.651 | 0.590 | 0.560 | 0.533 |
| Hagai2018_pig-lps | 0.543 | 0.483 | 0.489 | 0.455 |
| Hagai2018_rabbit-lps | 0.453 | 0.385 | 0.447 | 0.335 |
| Hagai2018_rat-lps | 0.666 | 0.521 | 0.453 | 0.453 |
| Hagai2018_rat-pic | 0.536 | 0.420 | 0.364 | 0.289 |
| Reyfman2020_alvmac<sup>c</sup> | 0.170 | 0.094 | 0.004 | 0.004 |
| Reyfman2020_pneumo<sup>c</sup> | 0.269 | 0.099 | 0.004 | 0.004 |

<sup>a</sup>Concordance with the ground-truth DEGs obtained with the bulk datasets using RankCompV3;
<sup>b</sup>Concordance with the ground-truth DEGs obtained with the bulk datasets using edgeR-LRT, the best method in ref [26];
<sup>c</sup>Concordance with the given bulk DEGs and no bulk profiles were provided.

Only in two datasets, Angelidis2019_alvmac and Reyfman2022_pneumo, did RankCompV3 perform poorly. The data quality of Angelidis2019_alvmac dataset was very poor and many genes were not detected in either the bulk or single-cell profiles. For the Reyfman2022 datasets, no bulk profiles were provided, and the given bulk DEGs were used as the ground-truth.

Furthermore, in comparison with the ground-truth obtained with edgeR-LRT, our results also show strong concordance. The median AUCC metric is 0.36 for the pseudo-bulk method and 0.29 for the single-cell method.

These results suggest that RankCompV3 can be effectively used for pseudo-bulk analysis, which can improve the performance of differential expression analysis in scRNA-seq data.

### Performance evaluation with the simulated single-cell dataset

Since we could not obtain all the truly differentially expressed genes in a real dataset, we simulated multimodal scRNA-seq profiles with scDD for cells under two conditions with 75 cells in each to test the performance. The dataset contains 2,000 differentially expressed genes (DEGs) equally distributed in four different modes and 18,000 non-DEGs equally distributed in two modes. The ROC curves are shown in Figure 3A. RankCompV3 has an AUC of 0.865, which is in the middle-tier of all 12 algorithms. The AUCs of Monocle2, SigEMD, Bimod, LR, MAST, DEsingle, Wilcoxon, edgeR, DESeq2, limma, and scDD are 0.649, 0.915, 0.820, 0.713, 0.620, 0.949, 0.632, 0.951, 0.966, 0.837, and 0.963, respectively.

**Figure 3.**
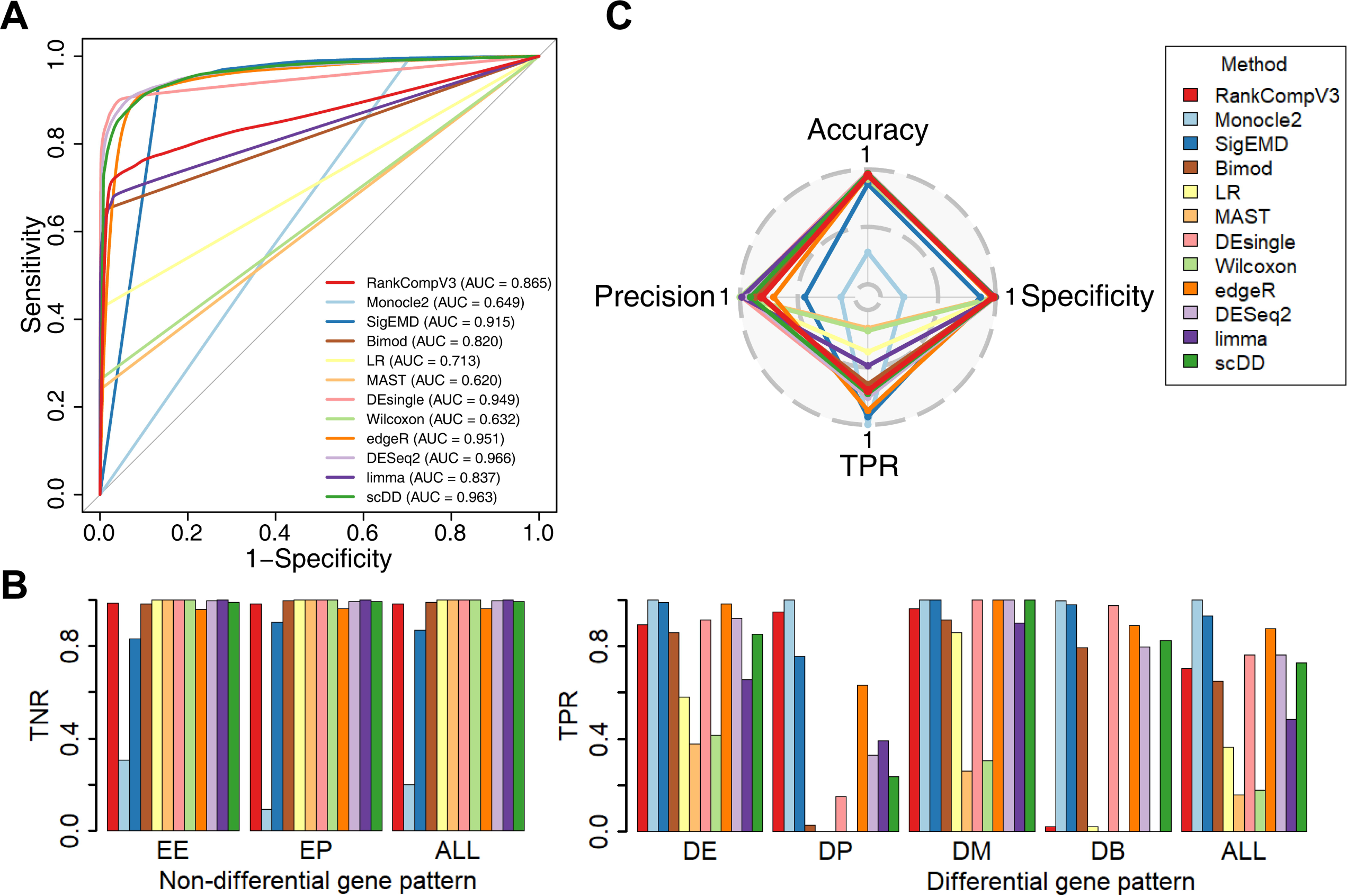
Simulated single-cell datase t. (A) ROC curves of RankCompV3 and other algorithms in the simulated scRNA-seq dataset. The area under each ROC curve (AUC) is shown in the legend. (B) TNRs and TPRs for the non-differential and differential genes with different expression modes in the simulated scRNA-seq dataset. (C) The radar map shows the Accuracy, Specificity, True positive rate and Precision of each algorithm.

The performances of the different algorithms are compared in Table 3 at a FDR < 0.05 threshold. RankCompV3 had a FPR of 0.018, a precision of 0.815, and an accuracy of 0.955, which were all better than or similar to the other algorithms. Compared to LR, MAST, and Wilcoxon, RankCompV3 showed a higher TPR while maintaining an extremely low FPR. Monocle2, SigEMD, and edgeR obtained extremely high TPRs, but they also contained a large number of false positives (FPs). Monocle2 had a TPR of 0.998, but its accuracy was only 0.122. This is because Monocle2 identified 16,383 DEGs among 20,000 genes, which introduced a larger FPR of 0.799 (Figure 3C).

**Table 3.**
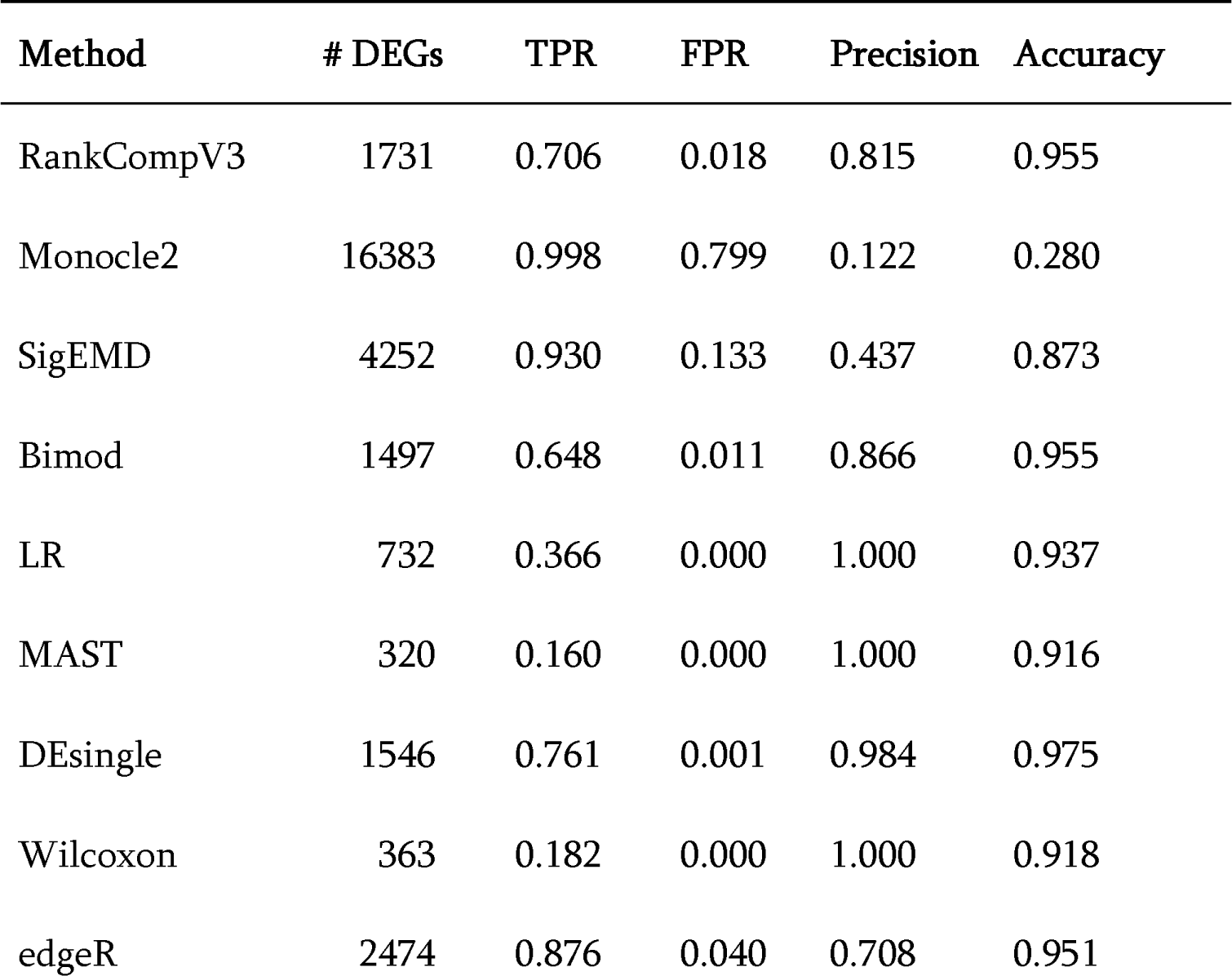
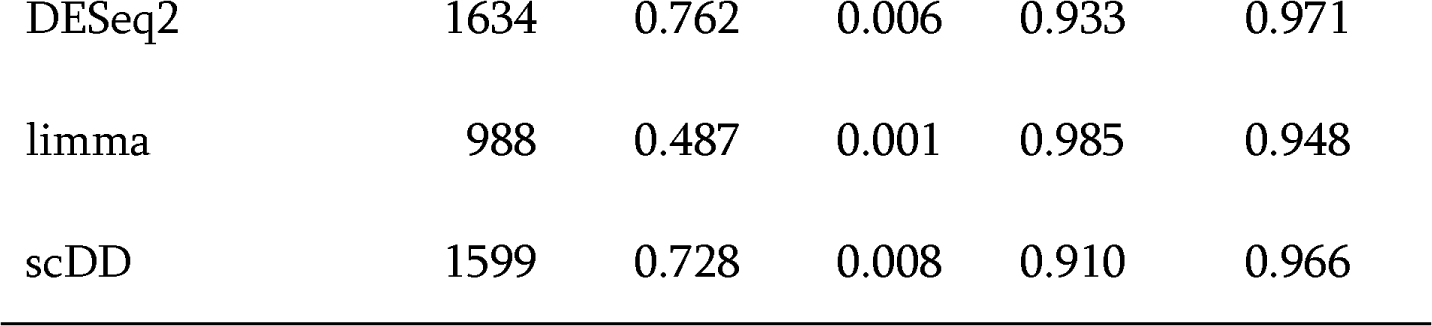
Area under the concordance curve (AUCC) for RankCompV3 for pseudo-bulk analysis scheme and single-cell analysis scheme in the ground-truth datasets from Squair et al. [26].

We evaluated the genes of different modes separately and compared the TNRs of non-DEGs of the two different modes and the TPRs of DEGs of the four different modes, as shown in Figure 3B. The results showed that the average TNR of the two modes of non-DEGs, EE and EP, was 0.982 using RankCompV3. The highest TPR of the four modes of DEGs, DD, DP, DB, and DM, was 0.962. For the DEG modes with no or low multimodality, DE and DM, the average TPR was 0.927. For the highly pleiotropic DEGs, DP and DB, the identification ability for DP was also very high (0.946), but it failed to detect DB genes. These results demonstrate that RankCompV3 strictly controls the FPR in the single-cell simulation dataset and has a good ability to detect DEGs of low multimodality and some DEGs with high multimodality.

In the simulated dataset, we randomly generated 10 pseudo-bulk profiles for each group to identify DEGs. The mean number of detected DEGs from the 10 random experiments was 1737.5 (standard deviation, SD = 13.3), and the mean number of true DEGs was 1413.3 (SD = 3.1). These numbers show a slight improvement compared to those obtained with the single-cell profiles directly (1731 and 1411, respectively). The mean AUCC metric was 0.884 (SD = 0.01), showing a strong concordance between the pseudo-bulk and the single-cell methods.

### Performance evaluation in real scRNA-seq dataset

Although the simulated dataset mimics numerous aspects of single-cell expression profiles, it cannot fully capture the complex characteristics of real data. To evaluate the performance of RankCompV3 on real data, we used the scRNA-seq dataset provided by Islam et al. [37], which consisted of the profiles of 48 mouse ES cells and 44 mouse embryonic fibroblasts. The top 1000 differential genes verified by RT-qPCR experiments were used as the gold standard DEGs (top1000).

Figure 4A shows the number of DEGs identified by each algorithm, the true number of DEGs (the number of genes that intersect with the top1000 genes), AUCC, and precision. RankCompV3 identified 1045 DEGs, of which 242 were true DEGs (TPR = 0.242). Although the number of true DEGs is the smallest, RankCompV3 has the highest precision and accuracy. Its AUCC concordance metric with the top1000 is lower than edgeR only (0.151 vs. 0.168).

**Figure 4.**
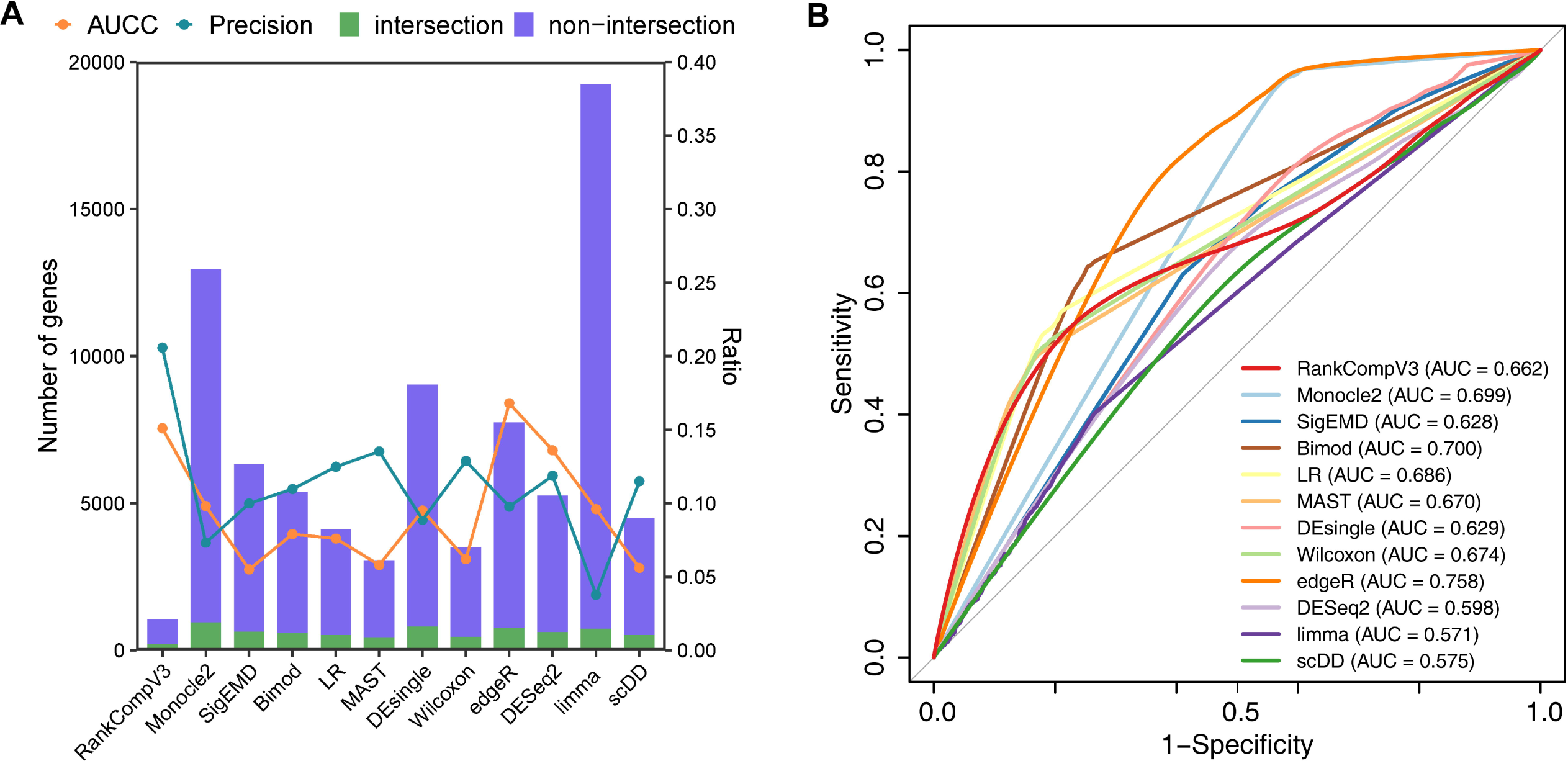
Real scRNA-seq dataset. (**A**) Performance in identifying DEGs in the scRNA-seq test dataset of GSE29087 where the top1000 genes confirmed by RT-qPCR were used as the ground-truth. The purple column represents the number of DEGs identified by each algorithm. The green area represents the number of intersection genes between the DEGs identified by the algorithm and the top1000 genes. The connected blue dots indicate TPRs and the connected orange lines indicate the area of the concordance curve (AUCC) metric. (**B**) ROC curves in the scRNA-seq positive test dataset of GSE29087. The top1000 genes were used as the ground-truth and the AUC values are shown in the legend.

RankCompV3 has a strictly conservative FPR (0.037), while maintaining a higher accuracy (0.932) than the other 11 methods which show high FPRs, ranging from 0.122 to 0.873. Among the other 11 algorithms, limma, Monocle2, DEsingle, and edgeR have the highest TPRs, which are 0.761, 0.931, 0.797, and 0.753, respectively. However, these algorithms also have high FPRs, ranging from 0.330 to 0.873. As a result, their accuracies are relatively low, ranging from 0.150 to 0.651.

The ROC curves are shown in Figure 4B. The AUC of RankCompV3 is 0.662, which is comparable to the other 11 algorithms. The AUCs of Monocle2, SigEMD, Bimod, LR, MAST, DEsingle, Wilcoxon, edgeR, DESeq2, limma, and scDD are 0.699, 0.628, 0.700, 0.686, 0.670, 0.629, 0.674, 0.758, 0.598, 0.571, and 0.575, respectively.

In conclusion, RankCompV3 can identify DEGs in scRNA-seq positive datasets. Compared to the high FPRs of many other methods, RankCompV3 may be more suitable for studies that require strict control of FPR.

### Effect of sample size to performance

To investigate the dependence of performance on sample size, we used the scRNA-seq dataset of LTHSC of two age-group mice provided by Kowalczyk et al. [39] to test the performance of RankCompV3. Subsets with sizes of 10, 30, 50, and 70 were randomly sampled for each age-group, and the sampling experiments were repeated 10 times. DEGs identified by the same algorithm in the entire dataset were taken as the gold standard.

As shown in Figure 5, the DEGs identified by RankCompV3 showed extremely low FPR in all subsets (< 0.04). As the sample size increased, the TPR of RankCompV3 increased whereas the FNR decreased. At a sample size of 10, the TPR and FNR of RankCompV3 were only lower than those of Monocle2 and sigEMD, and only lower than Monocle2 when the sample size was 30. It is worth noting that Monocle2 identified 12,759 out of 23,366 genes as DEGs in the case of the full dataset. These DEGs were used as the gold standard in the analysis of the subsets. This inevitably led to significantly superior TPR and FNR on one hand, but extremely high FPR and low accuracy on the other hand.

**Figure 5.**
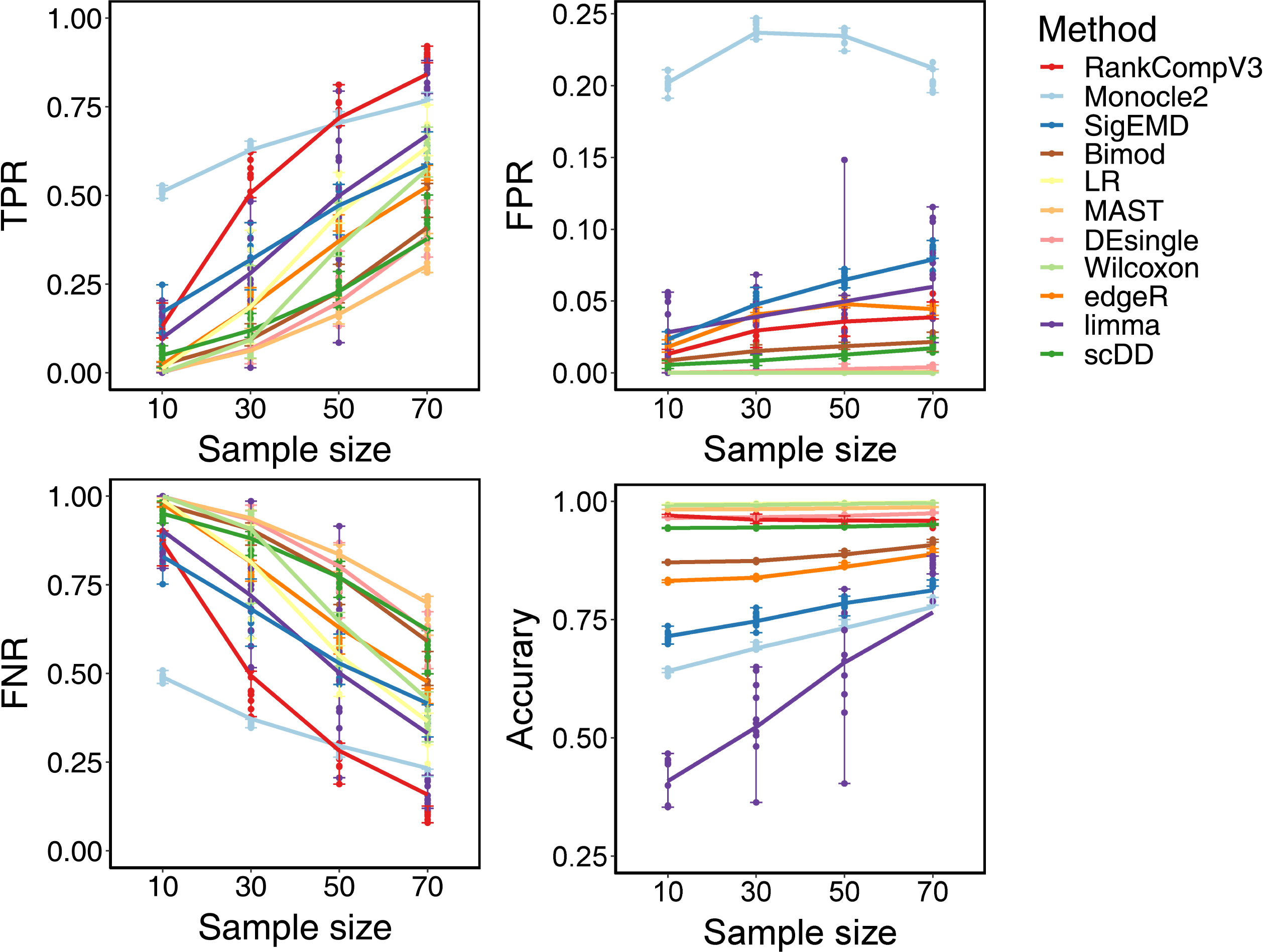
Performance dependence on sample size. The DEGs identified in the full dataset by each algorithm were used the ground-truth for the corresponding algorithm.

### Application in weak-signal datasets

In Misharin et al.’s study [44], they found that the differential expression of Siglecf can reliably distinguish Mo-AMs from TR-AMs in the bleomycin-induced early fibrosis. However, after 10 months of treatment with bleomycin, TR-AMs and Mo-AMs expressed similar levels of Siglecf and could not be distinguished by flow cytometry [40]. Only 330 DEGs were identified by two-way ANOVA (FDR < 0.05).

We compared the performance of our algorithm with that of other methods on this dataset. With FDR < 0.05, RankCompV3 identified 5023 DEGs. Monocle2, SigEMD, Bimod, LR, MAST, DEsingle, Wilcoxon, edgeR, DESeq2, limma, and scDD detected 1456, 2688, 182, 0, 3, 3, 0, 1224, 111, 79, and 720 DEGs, respectively. Several algorithms failed to detect or detected very few DEGs.

RankCompV3 identified more DEGs than the other methods, and its performance was not affected by the similarity of expression levels of Siglecf in TR-AMs and Mo-AMs. These results suggest that RankCompV3 is a more robust method for identifying DEGs in scRNA-seq data.

Through functional analysis, we confirmed that many DEGs identified by RankCompV3 have been shown to be associated with the differentiation of Mo-AMs and TR-AMs and the development of pulmonary fibrosis. For example, Siglecf can reliably distinguish Mo-AMs from TR-AMs [44]. Vcam-1 is a TGF-β1 responsive mediator that is upregulated in idiopathic pulmonary fibrosis [45]. SPARC drives pathological responses in non-small cell lung cancer and idiopathic pulmonary fibrosis by promoting microvascular remodeling and the excessive deposition of ECM proteins [46]. FGF2 inhibits pulmonary fibrosis through FGFR1 receptor action [47]. Adam8 deficiency increases CS-induced pulmonary fibrosis [48]. Macrophages expressing SPP1 proliferate during pulmonary fibrosis [49].

Six representative genes are shown in Figure 6. The first three genes, Sparc, Fgfr1, and Adam8, were DEGs identified by RankCompV3 only, whereas the other three, Ctnnb1, Anxa5, and Cap1, were DEGs identified by edgeR, limma, Monocle2, and SigEMD, but not by RankCompV3.

**Figure 6.**
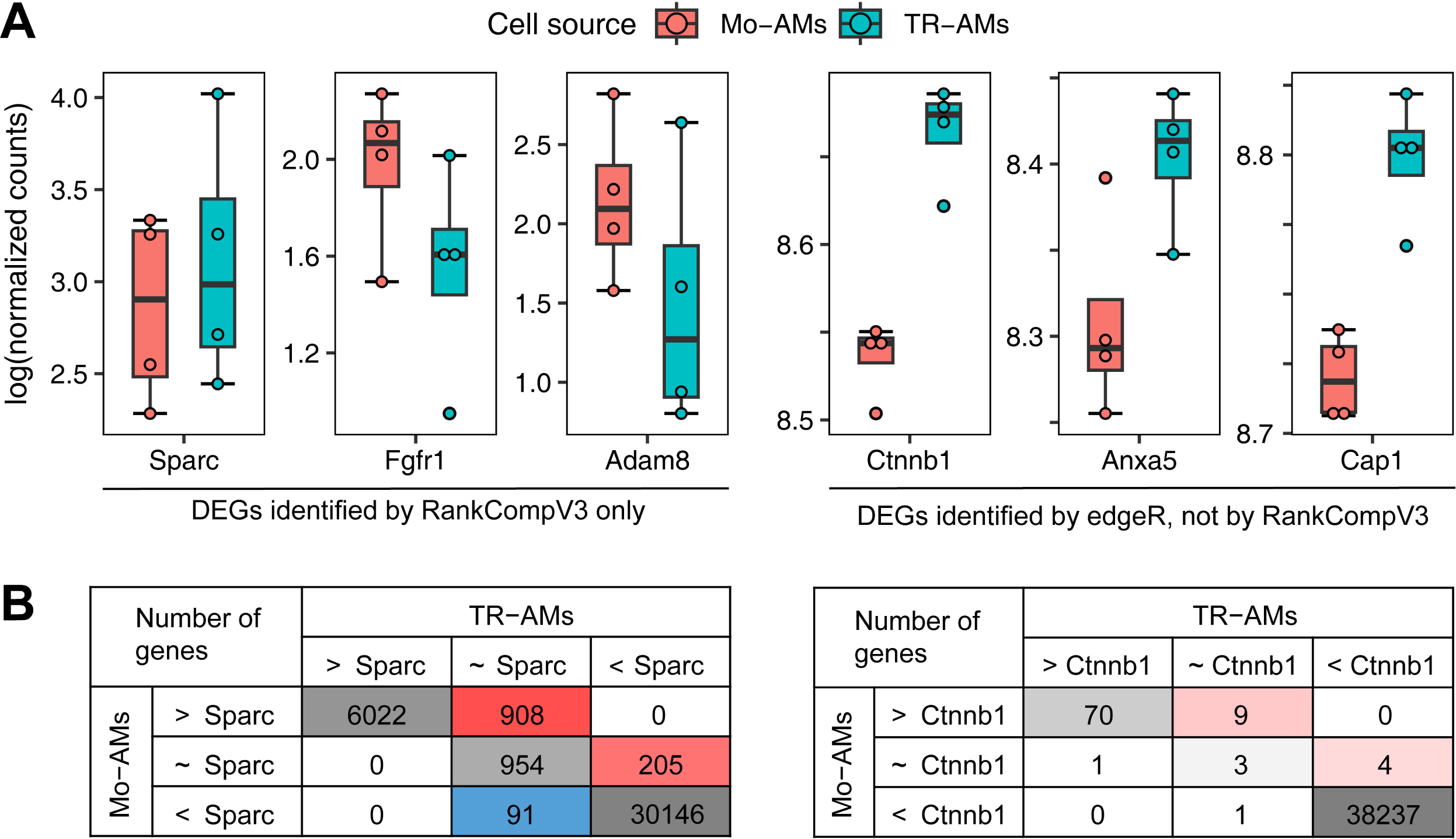
Box-whisker plots of the expression levels (log2 count) of 6 representative genes (A) and contingency tables for two of them (B). The first 3 genes (Sparc, Fgfr1, and Adam8) are DEGs identified by RankCompV3 only and the other 3 were non-DEGs by RankCompV3 but identified as DEGs by edgeR, limma, Monocle2, and SigEMD. Red cells in the contingency tables support that the gene is upregulated in the treatment group (TR-AMs), and blue cells support downregulation. The color intensity is proportional to the logarithm of the number of genes in each cell plus 1.

To demonstrate why the former are considered as DEGs and the latter as non-DEGs by RankCompV3, we also tabulate the contingency tables for Sparc and Ctnnb1 in Figure 6. A large number of reference genes support that Sparc is upregulated in TR-AMs, although the expression levels are approximately the same in the two cell types. The evidence is not strong to support that Ctnnb1 is upregulated, although the difference in the expression levels is significant.

We can see that RankCompV3 is able to identify lowly expressed genes and is not biased towards highly expressed ones, which is a common problem for many single-cell DE methods [26]. It is worth noting that many other algorithms fail to detect any of these DEGs, and only Monocle2 (Siglecf, Spp1, and Vcam1), SigEMD (Spp1), edgeR (Siglecf, Spp1, and Vcam1), and DESeq2 (Spp1) identified 1 to 3 of the above functional-meaningful genes.

We performed pathway enrichment analysis (FDR < 0.05) for the DEGs identified by RankCompV3. The DEGs were enriched in 82 pathways, of which five were associated with pulmonary fibrosis (Figure 7). Similarly, we performed pathway enrichment analysis on the DEGs identified by the other algorithms. Only five algorithms, Monocle2, SigEMD, DEsingle, edgeR, and DESeq2, recognized 1 to 3 of these functional pathways. Even when the significance threshold was relaxed to FDR < 0.2, the DEGs identified by Bimod and MAST were still not enriched in any of these pathways.

**Figure 7.**
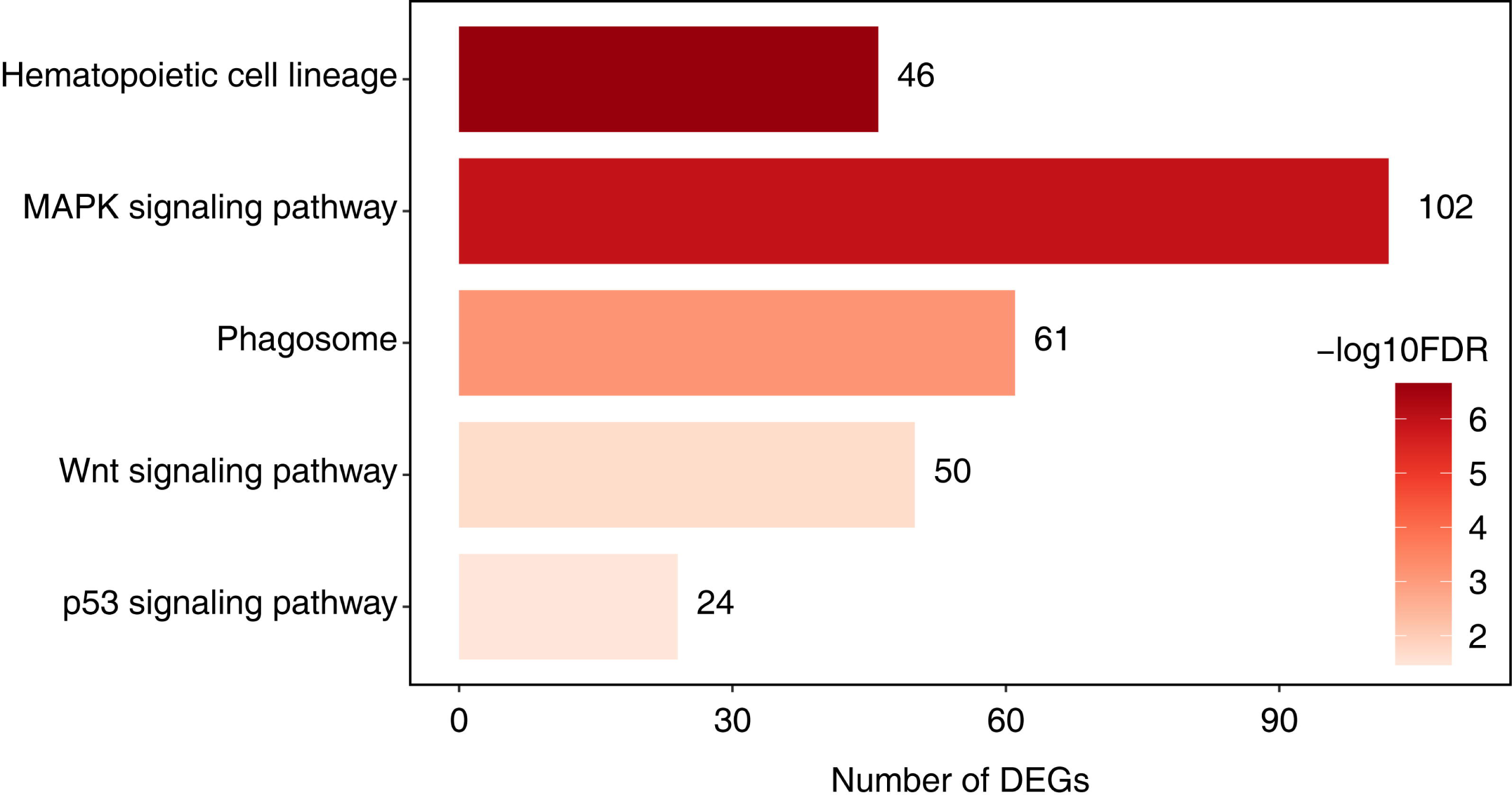
Pulmonary fibrosis-related pathways enriched with DEGs identified by RankCompV3. The x-axis value (also shown next to the bars) indicates the number of DEGs and color indicates the FDR value of the enriched pathway.

The Wnt signaling pathway was specifically identified by RankCompV3. The Wnt signaling pathway is an important pathway promoting pulmonary fibrosis [50], and studies have shown that targeting the Wnt pathway is a potential new treatment option for fibrosis [51].

In conclusion, RankCompV3 can detect weak biological signals that are functionally meaningful.

### Implementation and Runtime Analysis

For computational efficiency reason, we implemented RankCompV3 in Julia, a modern scientific computing language that is both easy to use and has performance on par with C [52]. The RankCompV3 package can be directly added using the Pkg.add(“RankCompV3”) function in Julia and it can also be called in R via julia_installed_package("RankCompV3"). The source code is available at https://github.com/pathint/RankCompV3.jl.

A typical analysis takes a few minutes, depending on the sample sizes (number of profiles in each group), number of genes, and number of execution threads. The most time-consuming step in RankCompV3 is the comparison of a large number of gene pairs. The time complexity of a naïve implementation is *O* (*nN* ^2^), where *N* is the number of genes passing the filtering step and *n* is the total sample size.

In Supplementary Table S2, we show the average runtimes of the 12 tools for the test in Figure 3. Even using a single thread, RankCompV3 is able to achieve faster or similar speed compared with other algorithms. Furthermore, the time cost can be significantly reduced by increasing the number of execution threads. In Supplementary Figure S2, we show the parallel runtimes for 1 to 8 threads on a single node.

## Discussion

Relative expression orderings (REOs) are stable in normal samples, but they are often disrupted in diseased samples [53]. This led us to develop two versions of RankComp [7], which can be used to identify DEGs at both the population and individual levels. These algorithms are insensitive to batch effects and have the advantage of being able to integrate datasets from different sources.

Fisher’s exact test was used in the original version of RankComp to calculate the significance level of the 2×2 contingency tables. RankCompV2 [8] added a filtering step to obtain a stable list of DEGs using non-DEGs as the background. This improvement circumvents the correlation effect between the true DEGs and their paired genes. But we failed to recognize that the REO analysis is a matched pairs design. The stability and direction of the REO of a gene-pair are interrelated in the two groups; they should not be counted independently.

In this study, we modified the contingency tables to tabulate the counts of 9 possible REO combinations for matched pairs design. McNemar’s test is designed for the 2×2 contingency table of a matched pairs experiment. The McNemar-Bowker test extends it to the variables of general *k* categories other than dichotomous variables and it is also called symmetry test of contingency tables. However, in this study, the three REO outcomes are ranked categories, and McCullagh’s method is a more appropriate choice. This method uses a logistic model to compare ordered categorical data in matched pairs experiments.

Previously, we showed that our RankComp algorithms based on REOs can be applied to microarray and bulk RNA-seq profiles and proteome profiles [11,12,54]. However, their applicability to scRNA-seq data has not been explored.

The scRNA-seq profiles tend to exhibit multi-mode expression patterns, heterogeneity, and sparsity compared to the bulk RNA-seq profiles. This makes it challenging to identify differentially expressed genes (DEGs) in scRNA-seq data. Many algorithms have been developed for scRNA-seq data to deal with dropouts or multi-mode patterns. However, these algorithms often cannot deal with both simultaneously. Additionally, many algorithms developed specifically for scRNA-seq have high FPRs and are affected by the number of cells and signal strength of the datasets. Algorithms developed for bulk RNA-seq data have also been shown to perform well in scRNA-seq data [27,28]. For example, limma, edgeR, and DESeq2 are all effective at identifying DEGs in scRNA-seq data.

In our study, we found that RankCompV3 exhibits an extremely low FPR in both simulated scRNA-seq data and a real negative dataset. Although DEseq2 and DEsingle perform well on the simulation dataset, but they have moderate or low performance on other datasets. This may indicate that they have a performance advantage in those datasets that satisfy an ideal distribution, but in real scenarios where the distribution is more complex, these methods often result in poorer performance. Additionally, RankCompV3 showed a lower FPR and higher accuracy than limma, edgeR, and DESeq2 in a positive test dataset of scRNA-seq.

Another advantage of the RankComp algorithms is that either the counts or normalized data (such as RPKM, FPKM, or TPM) can be used. As a heuristic method, RankCompV3 does not rely on a particular distribution of the gene profiles. The implication of normalization to DEG identification was discussed in a previous work [7].

Pseudo-bulking is a method that has been found to be effective in improving differential expression analysis in scRNA-seq data [26]. In pseudo-bulking, cells of the same type within a biological replicate are aggregated to a pseudo-bulk profile. This helps to lower the impact of dropouts in scRNA-seq, which are common due to the low sequencing depth of single cells.

Using the bulk profiles as the ground truth, pseudo-bulk analysis schemes can capture more true lowly expressed DEGs while lowering false positive with high expression. We have also implemented this method in RankCompV3. Tests with simulated and ground-truth datasets show that our method produces concordant results using either single-cell or pseudo-bulk methods. The pseudo-bulk method slightly improves the performance compared to the single-cell method. The results in Squair et al.’s show that the performance of edgeR-LRT is better than 13 algorithms, especially the results of pseudo-bulk analysis are better than those of single-cell analysis. Through our previous analysis, edgeR also maintains better performance advantages than other 10 algorithms in multiple datasets. Therefore, we conducted comparative tests against edgeR-LRT using single-cell and pseudo-bulk data, and the results showed that our algorithm RankCompV3 performed better than edgeR.

In RankCompV3, the quantitative expression level is not used. This means that the method is not biased towards highly expressed genes. The advantage of denser read counts, which leads to a more accurate ordering of genes in the pseudo-bulk profiles, is largely compensated by the larger number of single-cell profiles through the binomial test of the stability of a REO in a group.

The influence of sample size on the performance of the evaluated methods was investigated. Most methods showed little improvement with increasing sample size. However, RankCompV3 showed a gradual increase in TPR with increasing sample size. In terms of TPR and FNR, RankCompV3 was second only to Monocle2. However, Monocle2 achieved its high TPR and FNR by including a large proportion of genes as DEGs in the full dataset. This resulted in extremely high FPR and low accuracy.

In contrast, RankCompV3 achieved strict FPR control while maintaining high precision and accuracy. This was possible because RankCompV3 does not rely on the quantitative expression level of genes. Instead, it uses a binomial test to assess the stability of REOs in single-cell profiles. This makes RankCompV3 more robust to dropouts and other technical artifacts, and allows it to identify DEGs with high accuracy even in small datasets.

For weak-signal datasets, some common algorithms cannot capture differential expression signals. For example, in the pulmonary fibrosis dataset, LR, MAST, DEsingle, and Wilcoxon algorithms failed to identify or identify very few DEGs. However, our algorithm identified more DEGs than the other 11 algorithms, and the DEGs are significantly enriched with pulmonary fibrosis-related pathways.

One reason for detecting a relatively high number of DEGs is due to the less stringent control of the REO stability when the sample size is very small. In the case of a sample size of 4, the probability of observing all identical REO outcomes, e.g., *a* < *b* in 4 profiles, is 6.25% using the binomial model, even if the two genes have the same expression levels. This implies that the off-diagonal elements of the contingency tables may be higher than expected with the preset significance threshold. This is a limitation of the method that might lead to high FPRs when the sample sizes are too small.

However, the functional study of the DEGs identified by RankCompV3 indicates that they are reliable in this weak signal dataset. This suggests that the algorithm is able to identify true DEGs even in the presence of noise and other technical artifacts.

In summary, the results of this study demonstrate that RankCompV3 is a promising algorithm for identifying DEGs in scRNA-seq data, even in small datasets with weak biological signals. It is able to achieve strict FPR control while maintaining high precision and accuracy, which makes it a valuable tool for identifying biologically relevant DEGs.

## Key Points

- RankCompV3 is a method for identifying differentially expressed genes (DEGs) in either bulk or single-cell RNA transcriptomics. It is based on the counts of relative expression orderings (REOs) of gene pairs in the two groups. The contingency tables are tested using McCullagh’s method.
- RankCompV3 has comparable or better performance than that of other conventional methods. It has been shown to be effective in identifying DEGs in both single-cell and pseudo-bulk profiles.
- Pseudo-bulk method is implemented in RankCompV3, which allows the method to achieve higher computational efficiency and improves the concordance with the bulk ground-truth.
- RankCompV3 is effective in identifying functionally relevant DEGs in weak-signal datasets. This is because the method is not biased towards highly expressed genes.

## Data Availability

The scRNA-seq simulation dataset was generated using the scDD package. Other datasets were downloaded from the Gene Expression Comprehensive Database (GEO, http://www.ncbi.nlm.nih.gov/geo/) and the accession codes are GSE82158, GSE54695, GSE29087 and GSE59114. The source code is publicly available at https://github.com/pathint/RankCompV3.jl.

## Contact

Corresponding author: Xianlong Wang, Department of Bioinformatics, Fujian Medical University.

First author: Jing Yan, Department of Bioinformatics, Fujian Medical University.

## Funding

XW was supported by Fujian Medical University (Grant No. XRCZX2017001) and the Natural Science Foundation of Fujian Province (Grant No. 2019J01294).

## Supplementary information

**Figure S1.** The distribution modes of DEGs and non-DEGs in the simulated dataset.

**Figure S2.** The running time of RankCompV3 versus the number of execution threads.

**Table S1.** The number of samples from bulk expression profiles and single-cell expression profiles were in the matched bulk and single-cell data.

**Table S2.** Average runtime of RankCompV3 and 11 tools.

## Notes

### Competing Interest Statement

The authors have declared no competing interest.

