## Supplementary_Figures for "RankCompV3: a differential expression analysis algorithm based on relative expression orderings and applications in single-cell RNA transcriptomics"

**Supplementary Materials**

**Simulated data types and distribution**

The 2,000 differentially expressed genes were evenly divided into four categories, including (1) DE genes have a unimodal distribution but different mean values in the two groups; (2) DP genes show a bimodal distribution in both the groups and have the same mean values, but the cell proportions are different in the two modes; (3) DM genes have a unimodal distribution in one group and a bimodal distribution in the other group, while one mode of the latter overlaps with the mode of the former; (4) DB genes also have a unimodal distribution in one group and a bimodal distribution in the other, but the mode of the former is located in the middle of the two modes of the latter. Among them, DE and DM have no or a low degree of multimodularity, while DP and DB have a higher degree of multimodularity. EP and EE are non-differentially expressed gene with a bimodal distribution and a unimodal distribution, respectively.

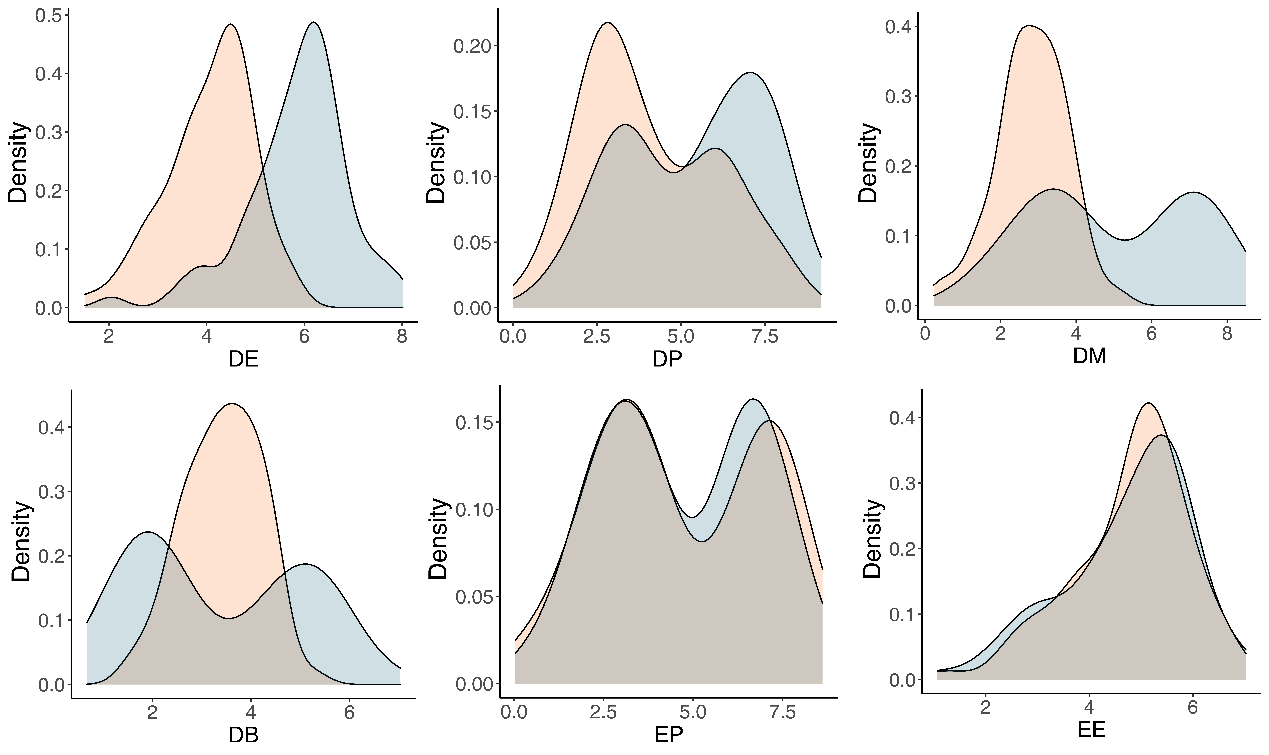

**Figure S1**. The distribution modes of DEGs and non-DEGs in the simulated dataset.

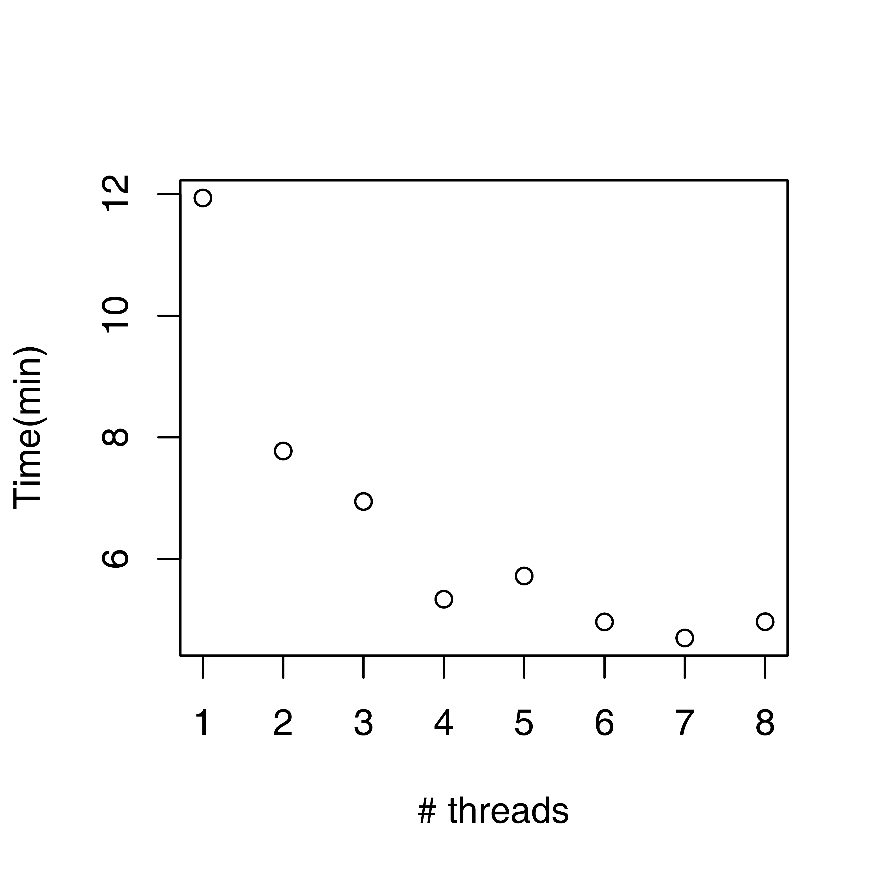

**Figure S2**. The running time of RankCompV3 versus the number of execution threads.
