## Supplementary_Table_S1 for "RankCompV3: a differential expression analysis algorithm based on relative expression orderings and applications in single-cell RNA transcriptomics"

**Table S1**. The number of samples from bulk expression profiles and single-cell expression profiles were in the matched bulk and single-cell data.

| **Datasets** | **Bulk** | | **Single-cell** | |
| --- | --- | --- | --- | --- |
|  | **Group1_num** | **Group2_num** | **Group1_cellnum** | **Group2_cellnum** |
| Angelidis2019_alvmac | 7 | 5 | 521 | 1183 |
| Angelidis2019_pneumo | 4 | 4 | 1190 | 3009 |
| CanoGamez2020_memory-iTreg | 6 | 6 | 3110 | 6131 |
| CanoGamez2020_memory-Th0 | 6 | 12 | 3110 | 4766 |
| CanoGamez2020_memory-Th2 | 6 | 6 | 3110 | 2893 |
| CanoGamez2020_memory-Th17 | 6 | 5 | 3110 | 5267 |
| CanoGamez2020_naive-iTreg | 6 | 6 | 2159 | 6588 |
| CanoGamez2020_naive-Th0 | 6 | 11 | 2159 | 2543 |
| CanoGamez2020_naive-Th2 | 6 | 7 | 2159 | 4040 |
| CanoGamez2020_naive-Th17 | 6 | 6 | 2159 | 5615 |
| Hagai2018_mouse-lps | 3 | 3 | 9195 | 8581 |
| Hagai2018_mouse-pic | 3 | 3 | 6515 | 8581 |
| Hagai2018_pig-lps | 3 | 3 | 6605 | 6148 |
| Hagai2018_rabbit-lps | 3 | 3 | 6650 | 10447 |
| Hagai2018_rat-lps | 3 | 3 | 5952 | 7325 |
| Hagai2018_rat-pic | 3 | 3 | 6127 | 7325 |
| CanoGamez2020_memory-Th1 | 6 | 6 | \ | \ |
| CanoGamez2020_naive-Th1 | 6 | 6 | \ | \ |
| Reyfman2020_alvmac | \ | \ | 13889 | 13684 |
| Reyfman2020_pneumo | \ | \ | 19736 | 6328 |
