## Supplementary_Table_S2 for "RankCompV3: a differential expression analysis algorithm based on relative expression orderings and applications in single-cell RNA transcriptomics"

**Table S2**. Average runtime of RankCompV3 and 11 tools.

| **Method** | **Language** | **Time (minutes)** |
| --- | --- | --- |
| RankCompV3 | Julia | 2.946413 |
| Monocle2 | R | 3.435453 |
| SigEMD | R | 6.357969 |
| Bimod | R | 0.36653 |
| LR | R | 1.067612 |
| MAST | R | 1.396969 |
| DEsingle | R | 29.9902 |
| Wilcoxon | R | 0.297589 |
| edgeR | R | 0.286038 |
| DESeq2 | R | 0.450563 |
| limma | R | 0.096508 |
| scDD | R | 47.24154 |
